# Alignment of multiple protein sequences without using amino acid frequencies

**DOI:** 10.1101/2024.06.05.597668

**Authors:** Veronika Shelyekhova, Roman Shirokov

## Abstract

Current algorithms for aligning protein sequences use substitutability scores that combine the probability to find an amino acid in a specific pair of amino acids and marginal probability to find this amino acid in any pair. However, the positional probability of finding the amino acid at a place in alignment is also conditional on the amino acids at the sequence itself. Content-dependent corrections overparameterize protein alignment models. Here, we propose an approach that is based on (dis)similarily measures, which do not use the marginal probability, and score only probabilities of finding amino acids in pairs. The dissimilarity scoring matrix endows a metric space on the set of aligned sequences. This allowed us to develop new heuristics. Our aligner does not use guide trees and treats all sequences uniformly. We suggest that such alignments that are done without explicit evolution-based modeling assumptions should be used for testing hypotheses about evolution of proteins (e.g., molecular phylogenetics).

## INTRODUCTION

Multiple sequence alignment (MSA) is a key step in many biological applications. It is difficult to compute [1, 2] and it is difficult to evaluate its accuracy [3]. In the last few years, however, there have been an upsurge in the number of algorithms that increase the accuracy for large (>10000) sets of sequences [4]. Most, if not all, current MSA algorithms maximize objective functions that are based on amino acid substitutability matrices, such as PAM [5] and BLOSUM [6]. The substitutability provides a statistical measure of how often an amino acid is replaced in evolutionary context. Substitutability combines the evolutionary, functional, and structural aspects of proteins. Therefore, it is fundamental for understanding sequence relationships that are beyond mere sequence identity, incorporating the biological significance of changes within sequences. Because substitution matrices penalize functionally unfavorable replacements, such as Alanine to Tryptophan, an aligner will try to put these residues into different sequence positions (columns). In addition, pairs of identical residues (one with two Alanine and another with two Tryptophan residues) score differently, with rare amino acids (Tryptophan) having a greater impact. It is relatively easy to move (mutate) one residue in a column with many Alanine residues. To do this in a column of Tryptophan residues is much costlier. This dependence of substitutability on “mutability” puts it at odds with measures of similarity [7]. However, the two terms are often used interchangeably in confusing ways.

Because the biochemical properties of amino acids, not their evolutionary past, define how proteins fold and function, we posit that sequence alignments, which reflect (and are evaluated by) the structural and functional properties of proteins, may be done using measures of similarity (or dissimilarity) between amino acids. Unlike substitution scores, similarity quantifications do not depend on the marginal probability to find an amino acid at a position.

We propose a transformation of a substitution matrix (BLOSUM in this work) that removes the information about the probability of finding an amino acid. Similar transformations can be done to obtain either similarity, or dissimilarity, scoring matrices. The resulting matrices differ only by the sign of their elements. We describe a global MSA scheme that is based on the Needleman– Wunsch algorithm [8]. It minimizes the total sum-of-pairs dissimilarity counts for all columns.

The dissimilarity measure sets a metric space on aligned sequences and, thus, allows comparing distances between them in a sound manner. In addition, the minimization has some algorithmic advantages over the commonly used maximization of substitutability. The informational contribution of dissimilar residues at regions of high similarity is large. However, when substitutability is maximized, a character standing in a column with many other identical among themselves characters might be ignored [9]. E.g., one Alanine may stay unnoticed among many Tryptophan residues in the column. Another advantage of the objective function minimization is that the search area of the dynamic programming matrix is inherently constrained for a given initial value of the function.

Most MSA algorithms reduce computational complexity by further exploiting evolutionary information. For example, progressive pairwise alignment relies on guide trees that cluster sequences using substitutability to access similarity. Topology of such guide trees strongly affects MSA quality [10, 11]. Conversely, evolutionary analyses might be biased when they are based on MSAs that use phylogeny-derived guide trees. Therefore, our alignment algorithm combines several heuristics that improve the result without entering the loop of the interrelationship between MSA and phylogeny.

Quality of our alignments was compared with those produced by two state-of-the-art algorithms: MAGUS/MAFFT [12, 13] and MUSCLE v5 [14]. It was done using two benchmark sets that differ in size of both query and reference alignments: QuanTest2 [15] and Balifam [14]. Our aligner performed as well as the other two on many of the QuanTest2 and Balifam query/reference subsets. However, it underperformed on those Balifam subsets that had low local similarity as defined by BLASTP [16] searches.

Thus, we show that a priori assumptions on the “true positional homology” in all columns [3] are not necessary for aligning proteins.

## RESULTS

### Dissimilarity scoring matrix and gaps

Values of the dissimilarity scoring matrix ***D*** were calculated as:

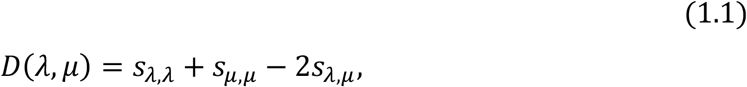

where: *D*(*λ*,*μ*) is the dissimilarity score for pair of amino acids *λ* and *μ*, and *s*_*λ*,*μ*_ are corresponding elements of BLOSUM matrix:

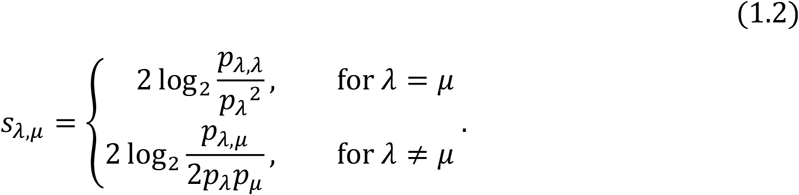

Here, *p*_*λ*,*μ*_ is the probability of two amino acids to replace each other; *p*_*λ*_ and *p*_*μ*_ are the probabilities of finding these amino acids. It follows that the *D*(*λ*,*μ*) values depend on the probability of observation of the pair (*p*_*λ*,*μ*_) but not on the background probabilities of finding the amino acids *λ* and *μ* at a position:

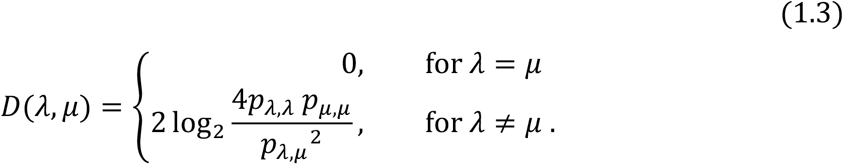

Unlike substitution matrices, the dissimilarity scoring matrix is hollow (*D*(*λ, λ*) = 0). This is equivalent to that pairs of identical amino acids do not contribute to the sum-of-pairs score.

Figure 1 shows the ***D*** matrix that is arranged to highlight its ability to relate the physicochemical properties of amino acids. Here, we start from the smallest value near the diagonal (2 for the Isoleucine-Valine pair). The matrix is then sorted so that the two upper diagonals, which are highlighted by thicker border lines, form the closest-follows-next path. Namely, the column for Leucine goes next after that for Valine because the Isoleucine-Leucine dissimilarity value (4) is the closest to that of the previous Isoleucine-Valine pair. The closest to the Valine-Leucine pair is the Valine-Methionine pair. So, the column for Methionine goes next, and so on:

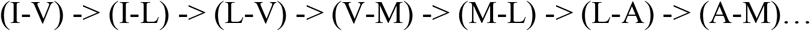

**Figure 1.**
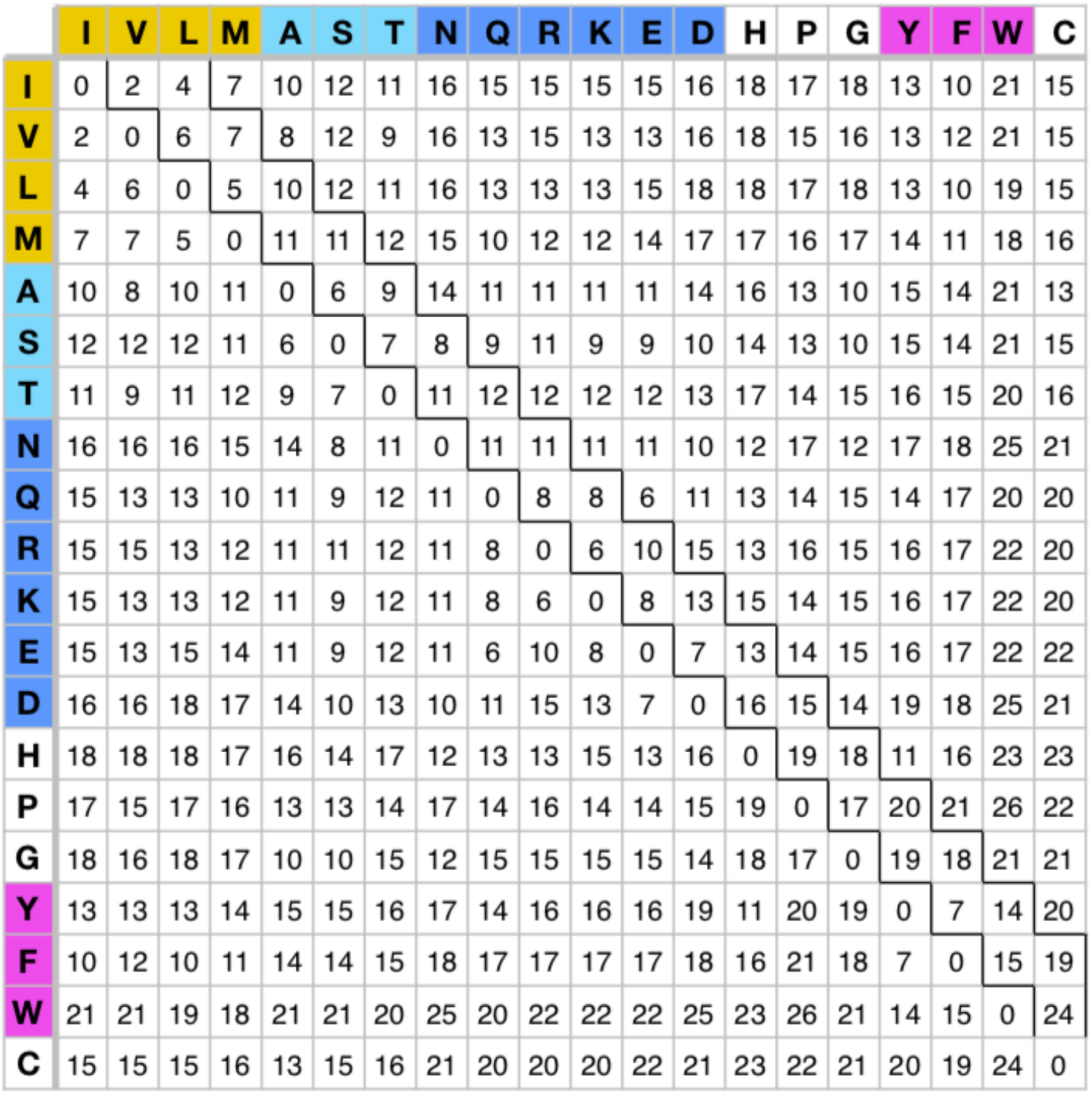
Matrix of dissimilarities between amino acids calculated from BLOSUM62. It is arranged to follow a path linking pairs with closes values of their dissimilarity scores.

There are multiple ways to sort the matrix as described. Such arrangements produce classifications of amino acids (highlighted by colored fill) that are like what was previously described [17].

Although the manipulation described by Eq. 1.1 removes the contribution of marginal probabilities, the dissimilarities calculated from the frequencies of amino acid pairs in blocks of aligned sequences might continue to reflect evolutionary selection preference. In other words, it is for evolution to decide the extent and importance of differences in physicochemical properties. To reveal that, we compared dissimilarity matrixes, which were calculated from BLOSUM matrixes for sequences with different degree of identity [18]. The BLOSUM45 matrix describes more divergent and, hence, evolutionary distant sequences. It is also implied that BLOSUM90 describes mostly evolutionary recent sequences. With increasing identity, the probabilities to find identical pairs are *p*_*λ*,*λ*_ expected to increase. This is reflected in that the diagonal elements increase in BLOSUM matrixes as the identity progresses. The dissimilarity scores also increase with the identity of sequences (Figure 3). The median value changed from 11 to 18. The increase in individual pairs was not uniform and it did not depend on the initial value at 45% identity. Dissimilarity score in some pairs changed by 3 or less (e.g., VI and CW, illustrated by blue lines). In other pairs, the dissimilarity increased by 10-11 points (yellow lines). Therefore, in some amino acid pairs the dissimilarity increased, or became more important, to a greater degree that in other. Such pairs are less frequently observed in alignment blocks with higher identity.

At first, we employed linear gap penalty, for which every space character in a gap gets the same dissimilarity cost *D*(*λ*, −) for any amino acid, and *D*(−, −) = 0. Later, we also developed the affine gap scheme that had different gap opening and extension costs. To our surprise, however, the affine gap heuristic did not improve the quality of alignment much (see: Alignment Evaluation). Using terminal gaps was helpful for certain benchmarking sets. For the dissimilarity matrix calculated from BLOSUM62, reasonable alignments could be done with the space character penalty in the range of 13-16, and 14 was the best. Accidentally, the mean value of the dissimilarity matrix elements for non-identical pairs is 14.5.

### Alignment method

For a given set of aligned sequences *s* ∈ ***S*** (∣ ***S*** *∣*= *N*) and the dissimilarity matrix ***D***, our algorithm minimizes the total sum-of-pairs ***S****P*:

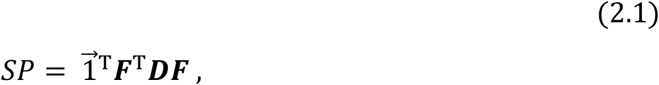

where frequency profile matrix ***F*** lists vectors of frequencies of characters in each position of alignment. The *SP* could be written as

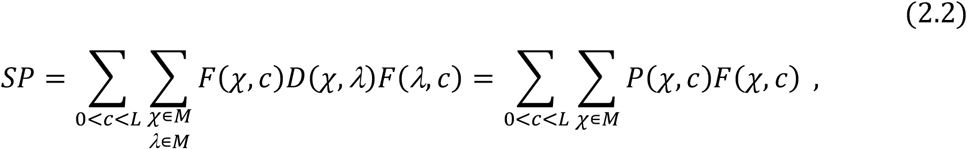

where: 𝒳 and ***A*** belong to the alphabet ***M*** consisting of 20 characters for amino acids and a special symbol for space (“−”); *c* is the index of *c*olumn in alignment of length *L*; *F*(𝒳, *c*) is an element of frequen*c*y profile ***F*** ; *P*(𝒳, *c*) is an element of profile *P* = ***D*** × ***F*** .

The *SP* may be indexed by sequences:

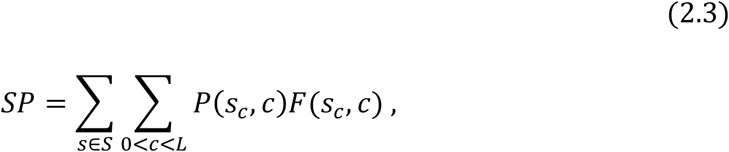

where *s*_*c*_ is a character in column *c* of sequence *s*.

If the space character element *D*(*λ*, −) *≥* 13, *D*(*λ, λ*) = 0, *D*(*λ, μ*) > 0, *D*(*λ, μ*) = *D*(*μ, λ*), *D*(*λ, μ*) + *D*(*μ, 𝒳*) ≥ *D*(*λ, 𝒳*) hold. Therefore, the dissimilarity matrix ***D*** endows a metric space on the alphabet ***M*** . It follows that the set of aligned sequences and the distance function *SP* also form a metric space Ω(***S***, *SP*). Note that adding gaps at different positions of a sequence creates different points in this space. The center point of an alignment is formally represented by the frequency profile ***F***. The inner sum in Equation 2.3, thus, has a meaning of the distance between sequence *s* and the central point.

A combination of greedy algorithms constructs global alignment by minimizing the *SP*. Our algorithms for profile-to-profile alignment (A1) and sequence alignment to a fixed profile (A2) are straightforward modifications of the Needleman–Wunsch dynamic programming procedure applied to protein profiles [16, 19-23].

An initial alignment was built from randomly shuffled sequences by profile merging (A1) that follows a balanced binary tree. To prevent a rapid growth of profile length, sequences in each group are realigned to the profile of the group (A2) and a new profile is produced before stepping to the next level of the tree. The alignment is refined by cycles of application of a fixed number of divide and merge steps based on A1, distortion by randomly varying the frequencies *F*(*𝒳, c*), and an optimization based on A2. The convergence is further improved by a divide-and-conquer approach that exploits the additivity of the *SP* by columns and minimizes it in a rolling window along the alignment (A3). Overall, the approach inherits all pros and cons of iterative descent algorithms for optimization in high dimensional space.

### Divide and Merge by Profile-to-Profile Alignment (A1)

To partition sequences, a “seed” sequence *s*_0_ ∈ ***S*** (a point in the metric space of aligned sequences) is chosen randomly. The *s*_)_ and the most distant from it point define the partition by closeness to them. The two profiles of the partition are aligned. If the divide-and-merge step does not reduce the *SP*, a new seed *s*_0_ is chosen so that it is most distant from previous *s*_0_ … *s*_*i−*1_ points. When the *SP* is improved, a new starting *s*_)_ is chosen randomly.

### Alignment to a Fixed Profile (A2)

If a point (sequence) moves in the Ω(***S***, *SP*) space toward the center point of the alignment (frequency profile ***F***), the sum of pairs *SP* decreases or remains the same (Eq. 2.3). Therefore, each sequence is aligned to profile ***F***. While gap insertions in the sequence are allowed, the profile is not altered. If the *SP* is reduced, columns consisting of only gaps are removed and a new profile is constructed. The fixing of the profile reduces the computational complexity of the algorithm from O(*Ll*) to O((*L* - *l*)*l*), where *l* is the length of un-gapped sequence. The removal of empty columns may increase the *SP* if the affine gap penalty cost is used. The problem is resolved by keeping some sequences unaltered to guarantee that every column has at least one amino acid character.

Profile ***F*** distortion is used to escape local minima of the target function. This is done by altering symbol frequencies and by inserting additional columns of gaps. Profile frequencies are changed according to the multinomial distribution with the number of trials equal to the number of sequences *N* and event probabilities being proportional to:

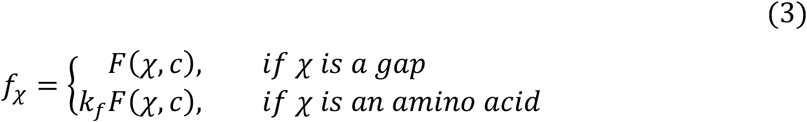

Here, a constant *k*_*f*_ > 0 determines the extent of distortion.

While frequency distortions are applied to all columns, insertions of aligned gap columns are done near columns with relatively large sum of pair in them:

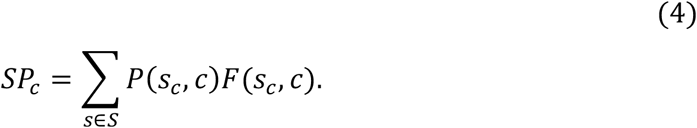

This is done according to the multinomial distribution with the number of trials equal to the number of columns to be entered and event probabilities that proportional to:

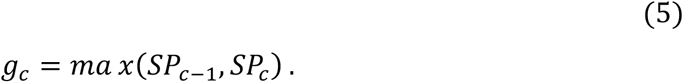

To stimulate the entering movements of symbols from the neighboring columns, the frequency for inserted columns of gaps *F*(−, *c*) is reduced to a new value 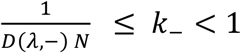.

The distortion parameters *k*_*f*_ and *k*___ that lead to a reduction in the *SP* are re-used to generate the new ones in next cycles. The insertion of columns of gaps are done only for the linear and terminal gap cost schemes, which do not change the *SP* with adding and/or removing gap columns. After distortions are applied, A2 is run repeatedly while the *SP* keeps decreasing, or a limiting number of iterations is reached.

### Rolling Window Alignment (A3)

The additivity of the *SP* by columns allows one to align separately a slice consisting of adjacent columns. If the slice is improved, it is spliced back to the alignment. The procedure is repeated along the length of alignment in a rolling window manner. The window sizes are chosen in the range of 2 to *L*/10 starting randomly and then are changed based on the performance at the previous cycle. The roll with a selected window size stops after a fixed number (2-3) of full-length rounds. In order to focus on regions that are not aligned well, the slices with their sum of pairs *SP*_*W*_ below the average 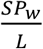 are skipped if 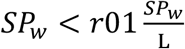, where *w* is the length of slice and *r*01 is a random number on the interval [0,1]. When terminal and affine gap cost schemes are used, the splicing of the slice may be affected by gaps outside it. Therefore, the slice boundary columns should not be used for the alignment.

### Alignment evaluation

We compared accuracy of our aligner with that of MAGUS/MAFFT[12, 13] and MUSCLEv5[14]. A detailed description and comparison of these and other current methods is given by Santus et.al [4]. We used QuanTest2[15] and Balifam[14] benchmarks. QuanTest2 is built by comparing the real structures of three reference sequences to the predicted secondary structure of test sequences. It has 151 test-reference sets with a thousand of sequences in each. Balifam is an expansion of BALiBASE3[24] obtained by addition of PFAM homologues. We chose the balifam10000 part, which has 36 test-reference sets with the number of reference sequences ranging from 4 to 142. After identical sequences were removed, the test Developer[25], scores using the qscore program[26]. To avoid the confusion with the sum of pairs objective function *SP*, we use the term Developer score.

First, the gap cost was optimized using the smaller QuanTest2 benchmark. Overall, the pair of 14 for the internal and 8 for the terminal gap gave maximal Developer scores. However, and to our big surprise, the affine gap cost scheme improved the Developer scores only by a few percent. Possibly, this is because the sequences were relatively short (the longest had 887 symbols and the median had 200).

Each point in Figure 3 represents a pair of Developer scores obtained by different aligners in 151 QuanTest2 datasets (protein families). Panel A compares our aligner (we call it “BB”) and MAGUS. Many points lay on the identity line. It means that our aligner performs on the corresponding datasets as well as MAGUS. Similarly, BB aligner is compared with MuscleV5 on panel B. The average Developer scores were somewhat higher for MUSCLE v5 (0.789 ± 0.016) and MAGUS (0.786 ± 0.015) than for BB (0.740 ± 0.018). However, BB was as good or even outperformed the competitors on many datasets. Panel C compares MAGUS and Muscle in the same manner.

The embedded benchmarks, which score only the alignment of reference sequences placed into the test, have multiple drawbacks [3]. One of them is that the test and reference sequences may poorly relate. It has been generally accepted that if two sequences have identity below 20%, the average alignment accuracy is not better than 50%[27]. Similar considerations should be applicable alignments on embedded benchmarks. Because Balifam has more reference sequences, we used it to analyze how the degree of relatedness of test and reference sequences affects the assessment of accuracy. Figure 4 compares Developer scores for MUSLE v5, MAGUS and BB on Balifam benchmark. Like in Figure 2, each dot corresponds to a protein family dataset. The coloring of dots will be explained below (Figure 5). Our aligner underperformed in about a third of all datasets.

**Figure 2.**
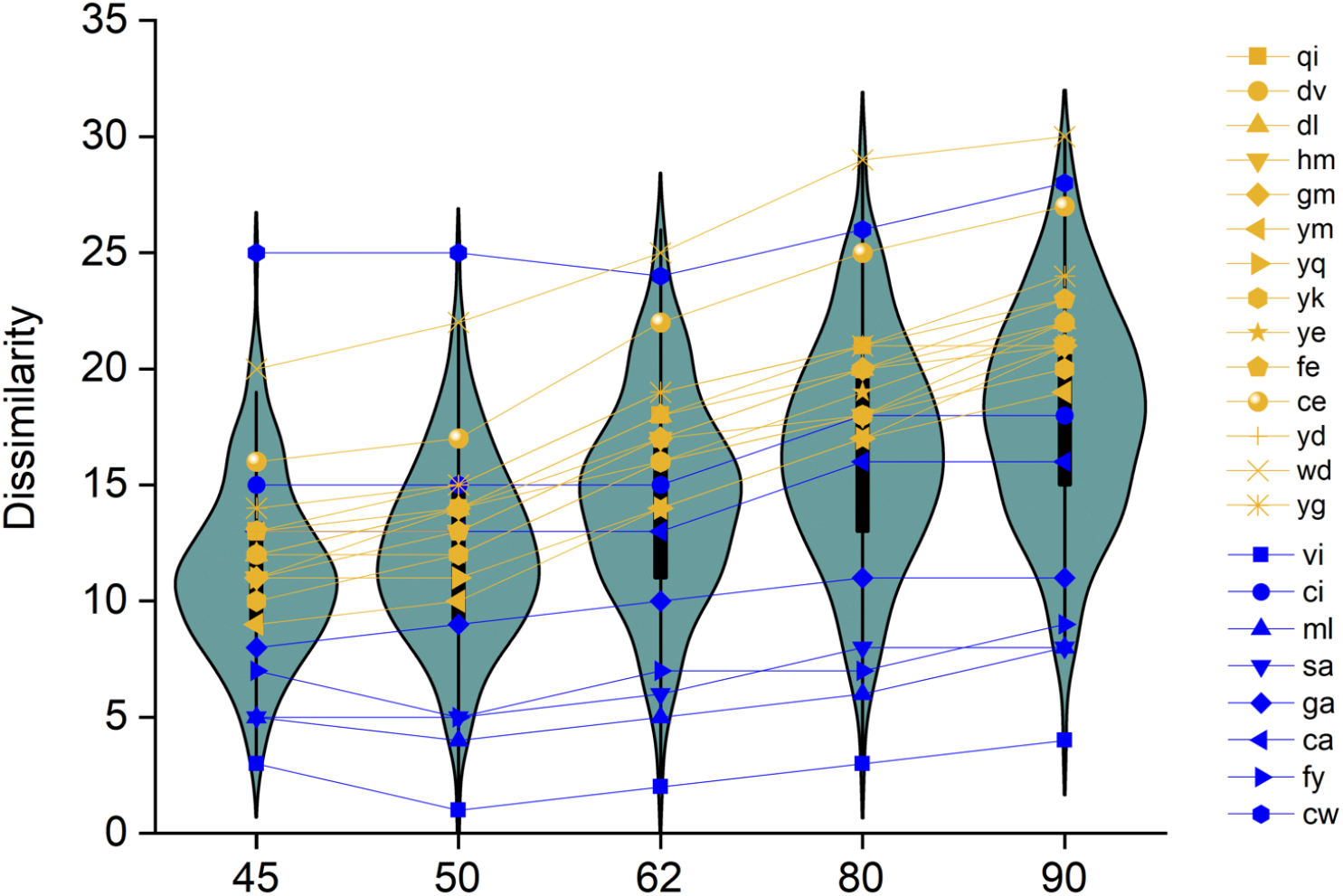
Changes in dissimilarity scores calculated from BLOSUM matrixes for different sequence identity. Violin plots illustrate distributions of all 190 pairs. Thick black lines show 25%-75% distribution interval, thin black lines show 1.5IRQ range. Blue symbols and lines show pairs that change dissimilarity by 3 and less when percent of identity changed from 45 to 90. Yellow symbols and lines show pairs that change dissimilarity 10 and greater.

**Figure 3.**
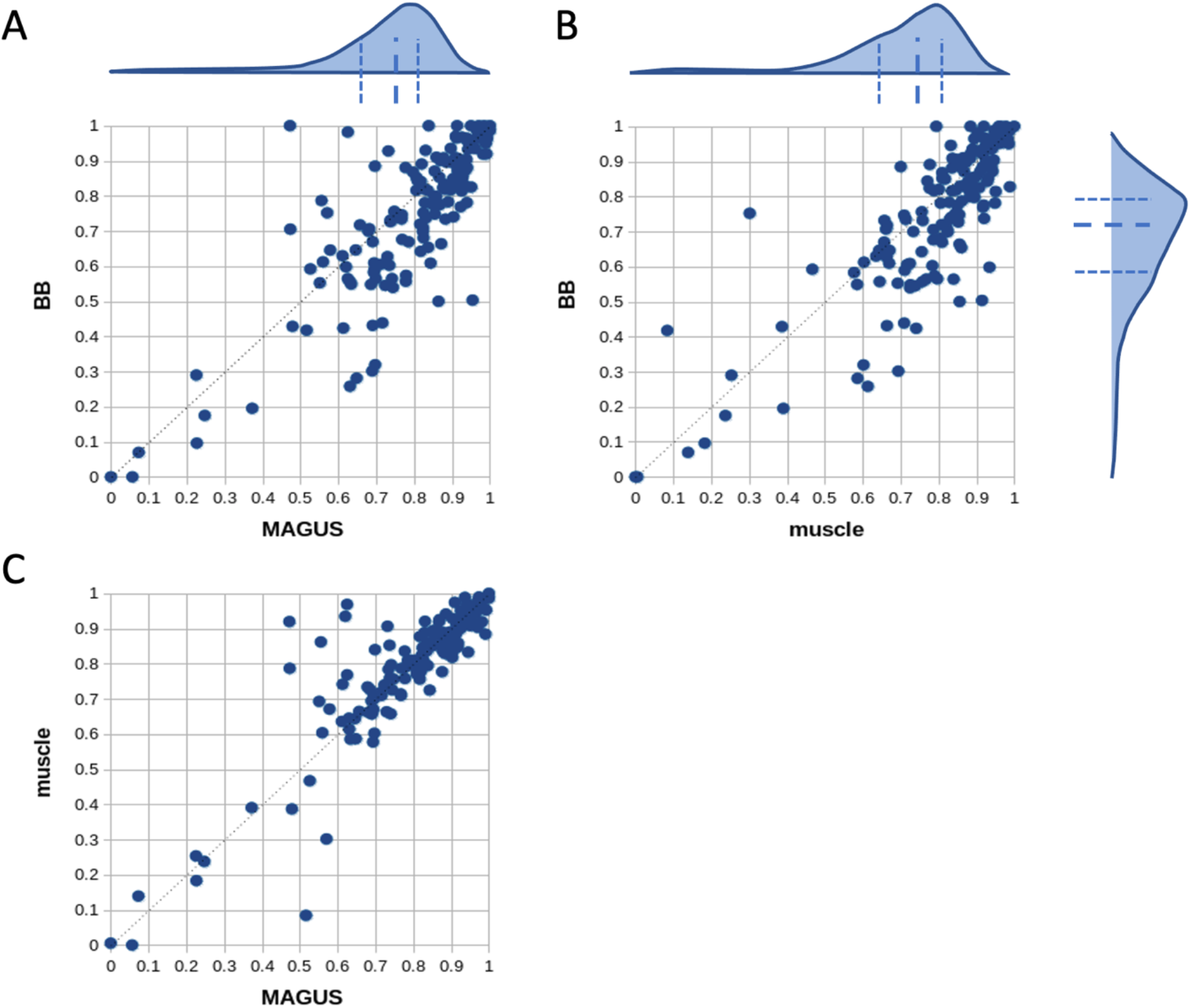
Comparison of Developer scores for BB, MAGUS and MUSCLE v5 aligners using QuanTest2 datasets. Each point shows Developer scores obtained by two aligners on one test/reference pair. The half-violin plots at the top of panel A and B illustrate distribution of Developer scores produced by MAGUS and MUSCLE v5 correspondingly. The half-violin plot on the right of panel B shows the distribution of Developer scores for BB aligner. The short-dashed lines show quartiles of the distribution. The thicker log-dashed lines show median values.

**Figure 4.**
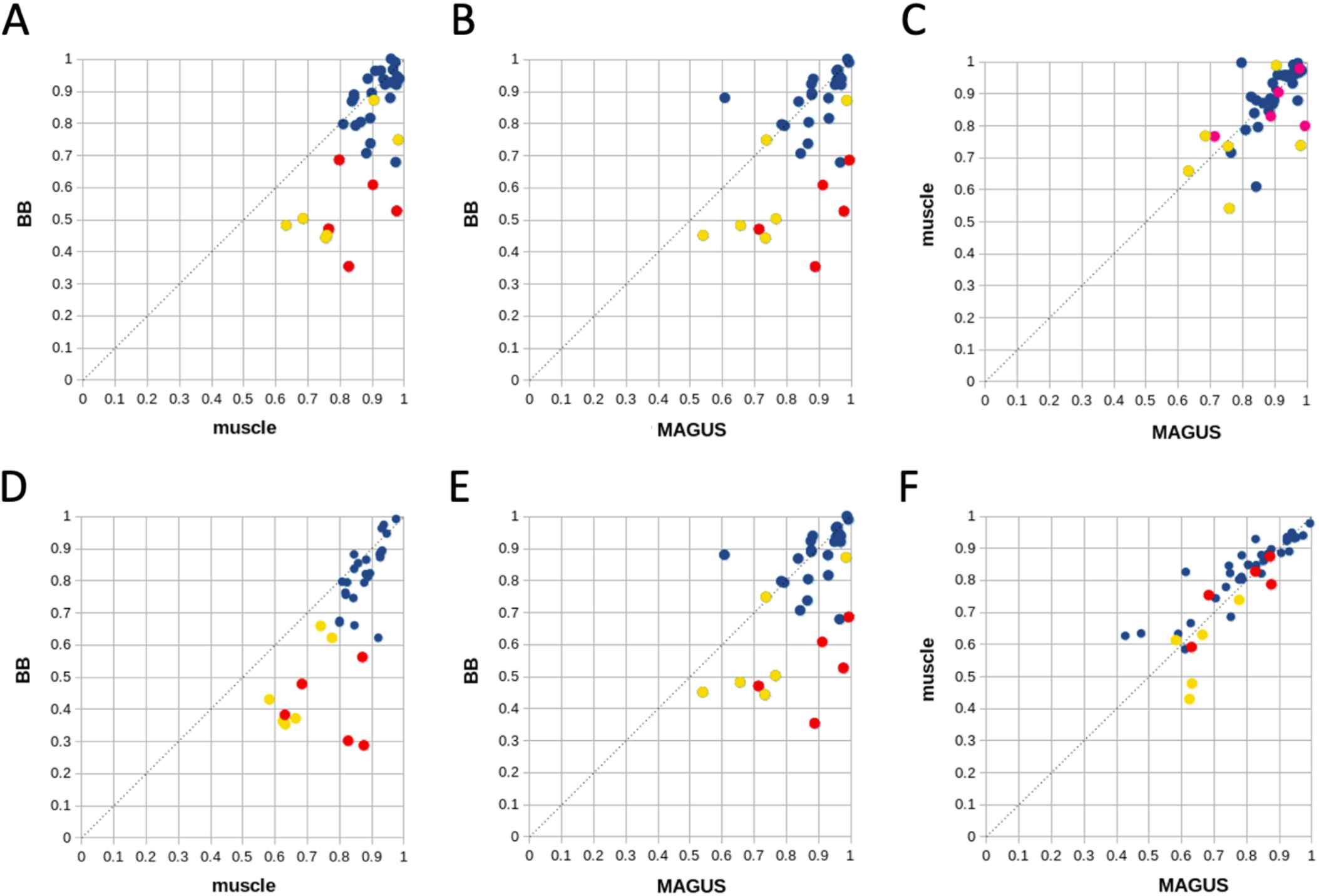
Comparison of Developer scores for BB, MAGUS and MUSCLE v5 aligners using Balifam benchmark datasets. Each point shows Developer scores obtained by two aligners on one test/reference pair. Panels A-C show Developer scores for columns that are selected as sites of significant structural identity. Panels D-F show the scores calculated for all columns. Colored dots correspond to datasets with low identity between test and reference sequences (See Figure 5).

**Figure 5.**
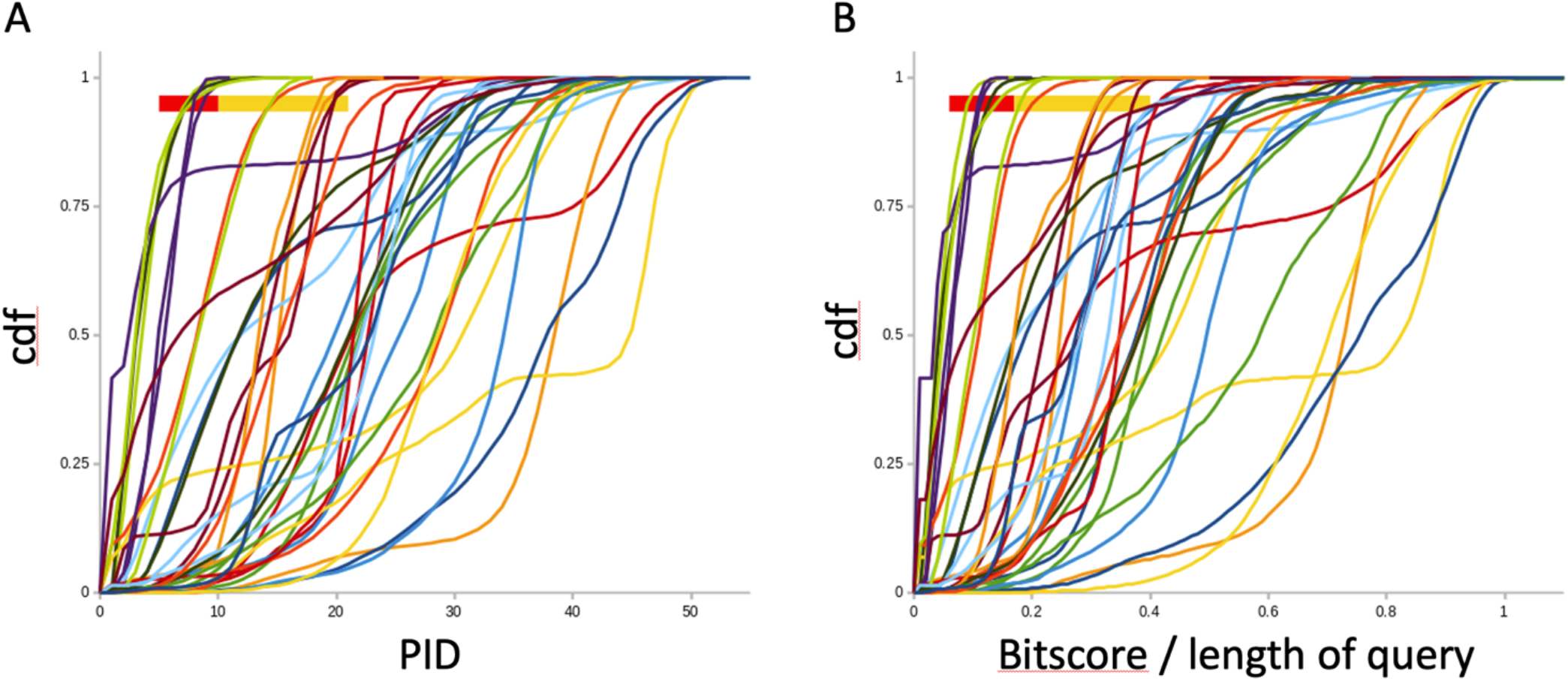
Cummulative distribution functions of percent of identity (Panel A) and the specific bitscore (Panel B) in Balifam datasets. The red/orange boxes at the top left corners of each graph show the 95% quantile. The datasets that cross the 95% quantile line at PID < 10 and bitscore/length < 0.15 are marked by red dots in Figure 4. Correspondingly, yellow dots on Figure 4 show datasets with PID < 20 and the specific bitscore < 0.4 at the 95% quantile.

Using blastp[16], high-scoring segment pairs (HSPs) between test and reference sequences in Balifam datasets were searched for. The maximal expectation value for a HSP was 0.05 in a test-reference pair. The threshold E-value was multiplied by the number of reference sequences when a test sequence was compared with the reference database. For each test sequence, the percent of identity (PID) with reference sequences and the total bitscore of the corresponding HSPs were obtained. The total bitscore was then divided by the test sequence length to obtain the specific bitscore.

Figure 5 shows cumulative distribution functions (cdfs) for PID (Panel A) and the specific bitscore (Panel B) in test sequences. Each smooth line corresponds to one Balifam test/reference pair. Some datasets have very little identity between test and reference sequences. Their cdfs saturated at small values of PID and the specific bitscore. The boxes at the top left corner of each panel mark the 95% quantile of the distributions. In the first five datasets, 95% of sequences have PID less than 10 (red box in Panel A). In the following six datasets, PID is less than 20 (yellow box). The same datasets also have low specific bitscores in HSPs (Panel B). Their corresponding specific bitscores at the 95% quantile were 0.15 and 0.4.

The HSPs bitscore could be converted to the corresponding BLOSUM62 score using *K* and *λ* parameters of the blastp runs. For a sequence with 100 amino acids, the specific bitscore of 0.2 corresponds to BLOSUM62 score of about 40, which is an average BLOSUM62 score for 6 identical residues. This is in accord with PID values estimated on the same group of sequences.

The benchmark datasets with low PID and the specific bitscores as determined by the procedure described in Figure 5 are shown by correspondingly colored dots in Figure 4. The red dots in Figure 4 show datasets with PID that is less than 5 in 95% of the test sequences. The yellow dots show the sets with PID(95%) that is less than 20.

Therefore, our aligner is not accurate, as judged by the Developer scores of embedded benchmarks when most of the test sequences are weakly related to the reference sequences. It is possible that both MAGUS and MUSCLE v5 scored better in these datasets because they align sequences in a progressive manner by combining pairs and/or clusters of sequences. This may result in that the final alignment is good only for the strongly related reference sequences. In contrast, our alignment method uniformly affects all sequences and does not rely on distance-related grouping.

## Conclusions

Content-dependent re-evaluation of substitutability and gap costs is one of many modeling assumptions of modern MSA algorithms. Amino acid function at a specific biochemical environment explains a big part of the apparent dependence of its availability and mutability on the local sequence content. The marginal probability to find an amino acid at a place is an essential constituent of substitutability counts. But it clearly is content-dependent. We show that reasonably good MSAs may be achieved by simply stacking up similar amino acids and they do not require accounting for mutability. Such MSAs that are based mostly on physicochemical properties of amino acids present less model-dependent datasets for testing hypotheses about evolution, such as for molecular phylogenetics. Analyses of MSAs that uniformly treat all sequences without the preconceived notion of sequence clusters are required for searching structures in the data. The overall averaging may appear to be disadvantageous when the sequences are known to be structured. However, the uniform alignment should be used for developing prior hypotheses.

The distance-based intuition is used in several heuristics that examine relationships between sequences. When amino acid (dis)similarities are considered, one can compare sequences by applying the notion of distance with the rigor of metric space.

## Acknowledgments

We would like to thank Hayk Hayotsyan for programming suggestions and Andy Harris for helpful discussion during the preparation of this work.

## Additional information

The program is available at: https://github.com/rshiroko/BB

